# On the consistency of duplication, loss, and deep coalescence gene tree parsimony costs under the multispecies coalescent

**DOI:** 10.64898/2026.02.20.707019

**Authors:** Nicolae Sapoval, Luay Nakhleh

## Abstract

Gene tree parsimony (GTP) is a common approach for efficient reconciliation of multiple discordant gene tree phylogenies for the inference of a single species tree. However, despite the popularity of GTP methods due to their low computational costs, prior work has shown that some commonly employed parsimony costs are statistically inconsistent under the multispecies coalescent process. Furthermore, a fine-grained analysis of the inconsistency has indicated potentially complementary behavior of duplication and deep coalescence costs for symmetric and asymmetric species trees. In this work, we prove inconsistency of GTP estimators for all linear combinations of duplication, loss, and deep coalescence scores. We also explore empirical implications of this result by evaluating inference results of several GTP cost schemes under varying levels of incomplete lineage sorting.

## 1 Introduction

Inference of the evolutionary history of a set of species—represented by a phylogenomic species tree—is challenging due to the discordance in the evolutionary histories of the individual genes within genomes [18,29]. Evolutionary histories of individual homologous sequences, modeled as gene trees [14], are often discordant with the species tree that traces the history of speciation events due to different biological processes. Gene duplication and loss [22,34] (GDL), and incomplete lineage sorting (ILS) [18] both contribute to this observed discordance. Statistical approaches for modeling these biological processes have emerged, with the multi-species coalescent (MSC) accounting for ILS [26,5,18] and DLCoal [27], MLMSC [10], and MSC-DL [20] accounting for GDL and ILS at the same time. In order to address the underlying biological question of phylogenomic species tree inference, multiple problem formulations and corresponding approaches have been proposed. In this work, we focus on summary methods that take as input a set of gene trees, and aim to reconstruct a species tree that optimizes some compatibility criterion.

Among the summary methods, there are multiple approaches that are proven to be statistically consistent under the MSC [1,21,9,13,12,11,19]. Also, more recent work has shown that some of the proposed approaches are also statistically consistent under DLCoal or variants thereof [25,16,8,23]. Conversely, some of the gene tree parsimony (GTP) methods that seek a species tree that minimizes the reconciliation cost with respect to the events of interest (duplications and losses [2,24], and deep coalescences [14,30,35]) have been previously shown to be inconsistent under the MSC [31]. However, in practice, GTP methods are still widely used due to easily interpretable optimization criteria, and relative computational efficiency, especially when compared to fully statistical methods that use maximum likelihood estimation or Bayesian inference. Furthermore, while the inconsistency of the individual GTP costs has been previously established, the results for joint optimization under combined criteria have not been theoretically analyzed [28].

In this work, we prove that any linear combination of the gene duplication and deep coalescence costs yields a GTP estimator that is inconsistent under MSC. In particular, we prove that there exists a species tree topology and a set of branch length parameters (i.e. an anomaly zone) for which the GTP estimator converges to an incorrect species tree topology. This is particularly interesting, since the anomaly zone for duplications arises from a symmetric species tree topology, and the one for deep coalescence score from an asymmetric topology. Additionally, we provide an extensive simulation-based comparison of different GTP scoring schemes which matches our theoretical results, and offers potential guidance for selecting optimal GTP scoring schemes in practical settings.

## 2 Preliminaries

### Definition 1

**(Gene tree parsimony estimator)**. *Given a collection of gene trees* 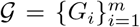 *and a cost function c*(*G, S*) *we call the species tree defined as*

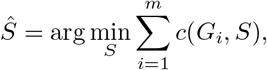

*the gene tree parsimony estimator of the species tree with cost function c*.

### Definition 2

**(Consistent estimator)**. *Let S*_T_ *denote the true species tree. Let* 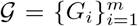 *be a set of gene trees arising from a multispecies coalescent (MSC) process on S*_T_. *Then, we say that a species tree estimator Ŝ*_*m*_ *is* consistent *if*

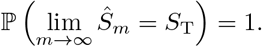

We define the gene duplication (*c*_*D*_(*G, S*)), gene loss (*c*_*L*_(*G, S*)), and deep coalescence (*c*_*X*_ (*G, S*)) costs analogously to [35]. Furthermore, analogously to [28] we define a generalized cost given by the linear combination of duplication, loss and deep coalescence costs

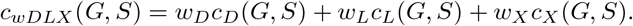

**Observation 1** *Let L*(*T*) *denote the multi-set of leaf labels of a tree T* . *If L*(*G*) = *L*(*S*) *and all labels are unique, then from Theorem 3*.*1 in [35] it follows that*

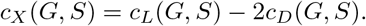

*Thus, it follows that*

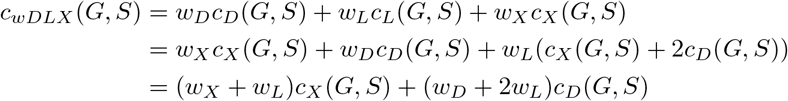

*for G satisfying the condition above*.

Thus, for the rest of the manuscript we will be concerned with proving statistical inconsistency of a cost given by *αc*_*D*_(*G, S*) + *βc*_*X*_ (*G, S*) where *α, β* ∈ ℝ.

**Observation 2** *Let* 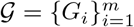 *be a set of gene trees arising from a MSC process on S*_T_, *and let* 𝒯_*N*_ *denote the set of all trees on N taxa, with S*_T_ ∈𝒯_*N*_ . *Let Ŝ*_*m*_ *be the gene tree parsimony estimator of the species tree with cost function c, and let* ℙ (*T*|*S*_T_) *denote the probability of observing gene tree T given species tree S*_T_ *under the MSC. From the strong law of large numbers it follows that*

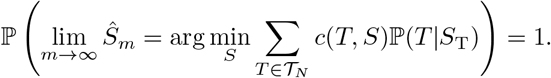

*Hence, it follows that the consistency of a gene tree parsimony estimator with cost function c can be determined by checking if* 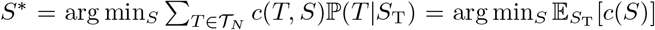 *is unique and equal to S*_T_.

## 3 Theoretical results

### 3.1 Statistical inconsistency of GTP under MSC

In prior work [31] it has been shown that the estimators under the deep coalescence cost are inconsistent when the true species tree topology is asymmetric. Furthermore, by explicitly analyzing all possible topologies for 4 species (Table 1), we note that the estimators under the duplication cost are inconsistent when the true species tree topology is symmetric.

**Table 1.** Probabilities, deep coalescence (*c*_*X*_) and duplication (*c*_*D*_) costs for each of the 15 rooted binary gene trees with leaf labels *A, B, C*, and *D* given either the species tree (*S*_1_; *λ*) = ((*a, b*) : *y*, (*c, d*) : *x*) or (*S*_4_; *λ*) = (((*a, b*) : *y, c*) : *x, d*).

| Gene tree $T_i$ | $\mathbb{P}(T_i\vert S_1; \lambda)$ | $c_X$ | $c_D$ | $\mathbb{P}(T_i\vert S_4; \lambda)$ | $c_X$ | $c_D$ |
| --- | --- | --- | --- | --- | --- | --- |
| $T_1 = ((a, b), (c, d))$ | $1 - \frac{2}{3}(e^{-x} + e^{-y}) + \frac{4}{9}e^{-(x+y)}$ | 0 | 0 | $\frac{1}{3}e^{-x} - \frac{1}{6}e^{-(x+y)} - \frac{1}{18}e^{-(3x+y)}$ | 1 | 1 |
| $T_2 = ((a, c), (b, d))$ | $\frac{1}{9}e^{-(x+y)}$ | 2 | 1 | $\frac{1}{6}e^{-(x+y)} - \frac{1}{18}e^{-(3x+y)}$ | 2 | 1 |
| $T_3 = ((a, d), (c, b))$ | $\frac{1}{9}e^{-(x+y)}$ | 2 | 1 | $\frac{1}{6}e^{-(x+y)} - \frac{1}{18}e^{-(3x+y)}$ | 2 | 1 |
| $T_4 = (((a, b), c), d)$ | $\frac{1}{3}e^{-x} - \frac{5}{18}e^{-(x+y)}$ | 1 | 1 | $1 - \frac{2}{3}(e^{-x} + e^{-y}) + \frac{1}{3}e^{-(x+y)} + \frac{1}{18}e^{-(3x+y)}$ | 0 | 0 |
| $T_5 = (((a, b), d), c)$ | $\frac{1}{3}e^{-x} - \frac{5}{18}e^{-(x+y)}$ | 1 | 1 | $\frac{1}{3}e^{-x} - \frac{1}{6}e^{-(x+y)} - \frac{1}{9}e^{-(3x+y)}$ | 1 | 1 |
| $T_6 = (((c, d), b), a)$ | $\frac{1}{3}e^{-y} - \frac{5}{18}e^{-(x+y)}$ | 1 | 1 | $\frac{1}{18}e^{-(3x+y)}$ | 3 | 2 |
| $T_7 = (((c, d), a), b)$ | $\frac{1}{3}e^{-y} - \frac{5}{18}e^{-(x+y)}$ | 1 | 1 | $\frac{1}{18}e^{-(3x+y)}$ | 3 | 2 |
| $T_8 = (((a, c), b), d)$ | $\frac{1}{18}e^{-(x+y)}$ | 2 | 2 | $\frac{1}{3}e^{-y} - \frac{1}{3}e^{-(x+y)} + \frac{1}{18}e^{-(3x+y)}$ | 1 | 1 |
| $T_9 = (((b, c), a), d)$ | $\frac{1}{18}e^{-(x+y)}$ | 2 | 2 | $\frac{1}{3}e^{-y} - \frac{1}{3}e^{-(x+y)} + \frac{1}{18}e^{-(3x+y)}$ | 1 | 1 |
| $T_{10} = (((a, d), b), c)$ | $\frac{1}{18}e^{-(x+y)}$ | 2 | 2 | $\frac{1}{18}e^{-(3x+y)}$ | 3 | 2 |
| $T_{11} = (((b, d), a), c)$ | $\frac{1}{18}e^{-(x+y)}$ | 2 | 2 | $\frac{1}{18}e^{-(3x+y)}$ | 3 | 2 |
| $T_{12} = (((a, c), d), b)$ | $\frac{1}{18}e^{-(x+y)}$ | 2 | 2 | $\frac{1}{6}e^{-(x+y)} - \frac{1}{9}e^{-(3x+y)}$ | 2 | 1 |
| $T_{13} = (((b, c), d), a)$ | $\frac{1}{18}e^{-(x+y)}$ | 2 | 2 | $\frac{1}{6}e^{-(x+y)} - \frac{1}{9}e^{-(3x+y)}$ | 2 | 1 |
| $T_{14} = (((a, d), c), b)$ | $\frac{1}{18}e^{-(x+y)}$ | 2 | 2 | $\frac{1}{18}e^{-(3x+y)}$ | 3 | 2 |
| $T_{15} = (((b, d), c), a)$ | $\frac{1}{18}e^{-(x+y)}$ | 2 | 2 | $\frac{1}{18}e^{-(3x+y)}$ | 3 | 2 |

This phenomenon is illustrated in Figure 1. The parsimony estimator using duplication score is inconsistent on a symmetric (with respect to the short branch) topology (Fig. 1A,C) and the parsimony estimator using deep coalescence score is inconsistent on an asymmetric topology (Fig. 1B,D).

**Fig. 1.**
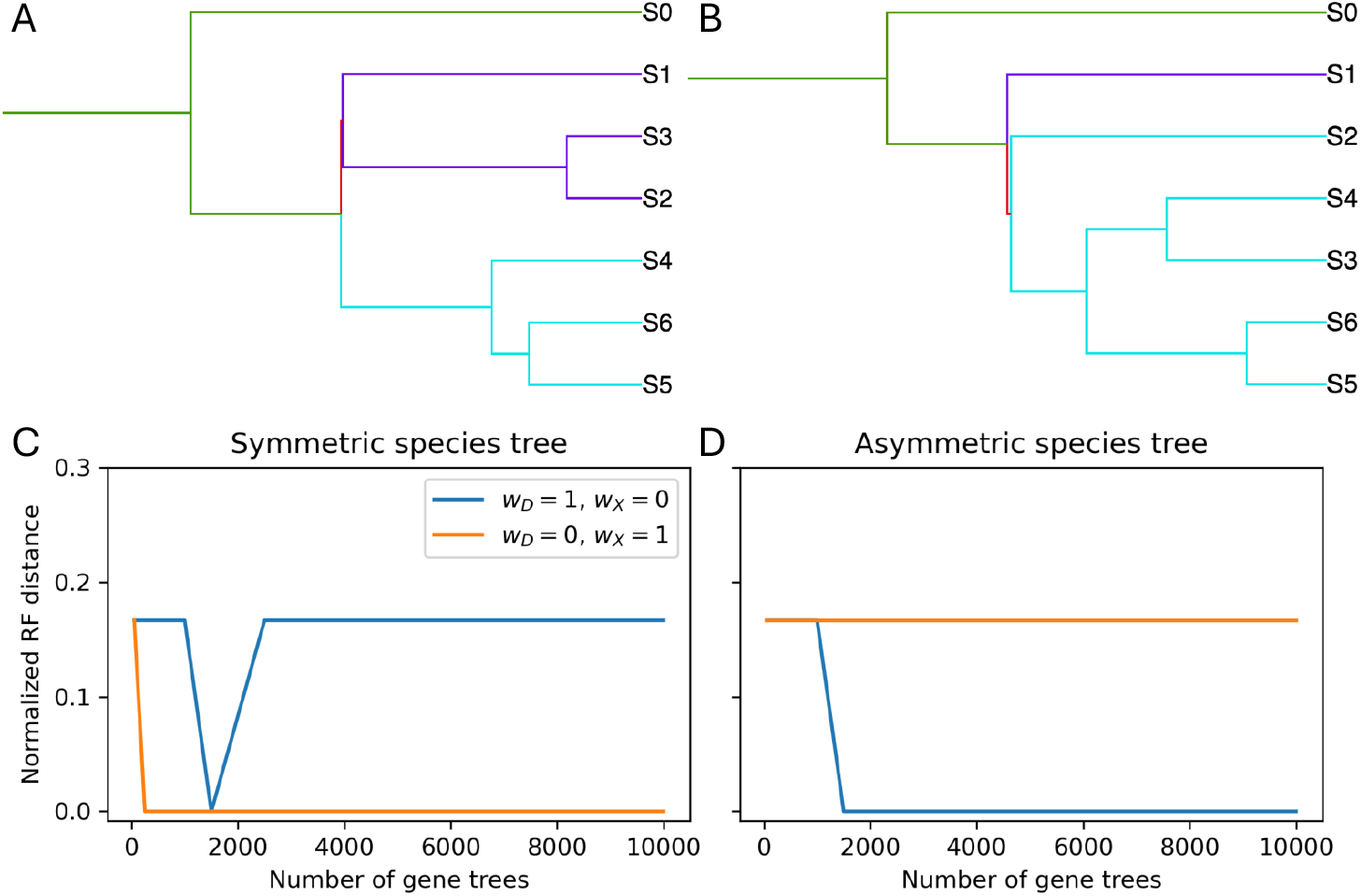
Duplication and deep coalescence costs are inconsistent on symmetric and asymmetric topologies, respectively. Two example species tree topologies, with one having a symmetric subtree structure associated with the ILS-driver branch (A), and another one with asymmetric subtree structure associated with the ILS-driver branch (B). (C, D) Normalized Robinson-Foulds distances between species trees inferred by GTP and the ground truth species tree as the functions of number of gene trees used. Blue line shows the duplication cost, and orange line shows the deep coalescence cost.

We can formalize and prove these observations as the following two lemmas.

#### Lemma 1.

*For all α* ∈ (0, ∞) *the gene tree parsimony estimator with the cost function αc*_*D*_(*G, S*) *is inconsistent for species trees with N* ≥ 4 *taxa under the MSC model*.

*Proof*. Let 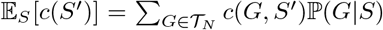 denote the expected cost of a species tree *S*′ for the ground truth species tree *S*. From Observation 2 it follows that a gene tree parsimony estimator with a cost function *c* is inconsistent under the MSC model if there exists *S*′ ∈ 𝒯_*N*_ s.t. 𝔼_*S*_[*c*(*S*′)] ≤ 𝔼_*S*_[*c*(*S*)].

Let *T*_1_ and *T*_4_ be the tree topologies as defined in the Table 1. We claim that for any *β* ∈ ℝ^∗^ there exist *x* ∈ (0, ∞) and *y* ∈ (0, ∞) s.t. for *S*_T_ = (*T*_1_; (*x, y*)) the following inequality holds

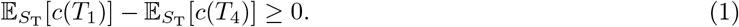

Using the probabilities and cost values listed in Table 1, we have

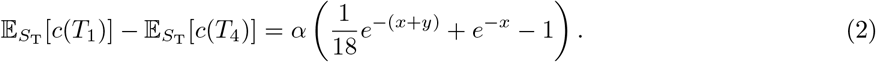

We seek values (*x, y*) s.t. the expression above is non-negative.

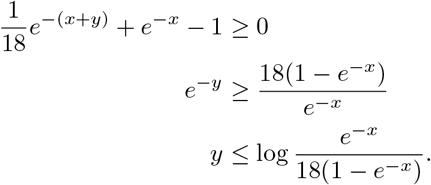

We note that for *x* < log(19*/*18) ≈ 0.054 the right hand side of the inequality is strictly positive. Thus, it follows that for any *x* ∈ (0, log(19*/*18)) and any 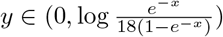 the estimator is inconsistent.

#### Lemma 2.

*For all β* ∈ (0, ∞) *the gene tree parsimony estimator with the cost function βc*_*X*_ (*G, S*) *is inconsistent for species trees with N* ≥ 4 *taxa under the MSC model*.

*Proof*. Than and Rosenberg [31] showed that the gene tree parsimony estimator with the cost function *c*_*X*_ (*G, S*) is inconsistent under the MSC model. By Observation 2, it follows that for *β* > 0 the species tree estimated under the cost *βc*_*X*_ is the same as the tree estimated under the cost *c*_*X*_.

In particular, Lemma 1 indicates that an estimator with the duplication cost favors asymmetric topologies in the anomaly zone for symmetric species trees, while Lemma 2 implies that an estimator optimizing the deep coalescence cost favors symmetric topologies in the anomaly zone for asymmetric species trees. Furthermore, we note that for any non-zero weight choice *β* for the deep coalescence cost the resulting combined cost will prefer symmetric topologies when the ground truth tree is asymmetric. More precisely, the following lemma holds.

#### Lemma 3.

*For any choice of weights α, β*∈ [0, ∞) *the gene tree parsimony estimator with the cost function c*(*G, S*) = *αc*_*D*_(*G, S*) + *βc*_*X*_ (*G, S*) *is statistically inconsistent for species trees with N* = 4 *taxa under the MSC model*.

*Proof*. The cases *β* = 0 and *α* = 0 follow from Lemmas 1, 2 respectively. Hence, we can assume that *β*≠ 0.

Let 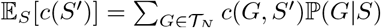 denote the expected cost of a species tree *S*′ for the ground truth species tree *S*. From Observation 2 it follows that a gene tree parsimony estimator with a cost function *c* is inconsistent under the MSC model if there exists *S*′ ∈ 𝒯_*N*_ s.t. 𝔼_*S*_[*c*(*S*′)] ≤ 𝔼_*S*_[*c*(*S*)].

Let *T*_1_ and *T*_4_ be the tree topologies as defined in the Table 1. We claim that for any *α* ∈ [0, ∞) and any *β* ∈ (0, ∞) s.t. *α/β* < 7 there exist *x* ∈ (0, ∞) and *y* ∈ (0, ∞) s.t. for *S*_T_ = (*T*_4_; (*x, y*)) the following inequality holds

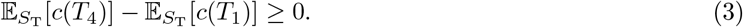

Using the probabilities and cost values listed in Table 1, we have

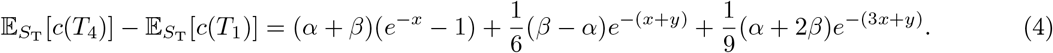

We seek values (*x, y*) s.t. the expression above is non-negative. We start by setting *y* = −log 0.9 0. ≈105 to obtain

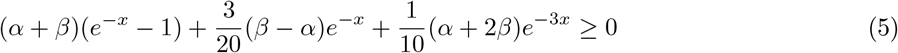

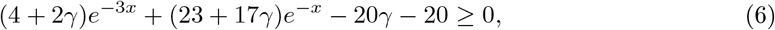

where *γ* = *α/β*, which is well defined since *β*≠ 0.

Now, let *f* (*x*) = (4 + 2*γ*)*e*^−3*x*^ + (23 + 17*γ*)*e*^−*x*^ − 20*γ* − 20, clearly *f* is a continuous function with lim_*x*→∞_ *f* (*x*) = −20*γ* − 20. Furthermore, we have that

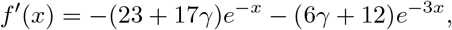

and hence the function *f* (*x*) is decreasing for any value of *γ* ∈ [0, ∞).

Thus, we know that *f* (*x*) is decreasing on [0, ∞) and *f* (0) = 7 −*γ* > 0 for *γ* < 7. Since lim_*x*→∞_ *f* (*x*) *<* 0 by the intermediate value theorem it follows that there exists some *x*_*r*_ ∈ (0, ∞) s.t. *f* (*x*_*r*_) = 0, and subsequently for all *x* ∈ (0, *x*_*r*_) we have *f* (*x*) *>* 0. Therefore, we can conclude that for *y* = −log 0.9 and *x* ∈ (0, *x*_*r*_) the inequality (3) holds for *γ* = *α/β* < 7.

For the case *γ* ≥ 7, we will consider *T*_1_ and *T*_4_ as before, and show that *α* ∈ [0, ∞) and any *β* ∈ (0, ∞) s.t. *α/β* ≥ 7 there exist *x* ∈ (0, ∞) and *y* ∈ (0, ∞) s.t. for *S*_T_ = (*T*_1_; (*x, y*)) the following inequality holds

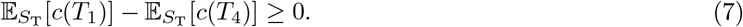

Using the probabilities and cost values listed in Table 1, we have

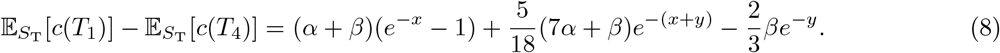

We seek values (*x, y*) s.t. the expression above is non-negative. We start by setting *y* =− log 0. ≈9 0.105 to obtain

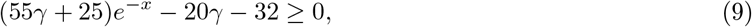

where *γ* = *α/β*. Proceeding the same way as in the previous case, we note that *f* ′(*x*) *<* 0 and lim_*x*→∞_ *f* (*x*) *<* 0. Furthermore, *f* (0) = 35*γ* − 7 *>* 0 for all *γ* ≥ 7. Hence, once again by the intermediate value theorem it follows that there exists *x* for which the inequality holds.

Thus, combining these two results, we can conclude that for any choice of *α, β* there exists a 4-taxon species tree for which the parsimony estimator is inconsistent.

Than and Rosenberg [31] provided a general framework for recognizing an embedded 4-taxon tree within a larger species tree, thus extending proofs of statistical inconsistency of parsimony methods from 4-taxon trees to all *N* ≥ 4.

#### Lemma 4.

*Let S*_T_ *be a species tree with 5 or more leaves, and let S*_1_ *and S*_4_ *be the four-leaf tree as in Table 1*. *Let* 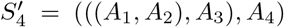 *denote the embedded asymmetric structure in S*_T_, *and let* 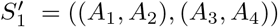 *denote an alternative symmetric structure (see [31], Section 3*.*3). Then the following inequalities hold:*

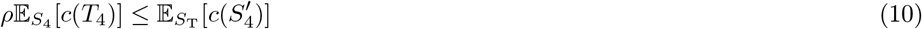

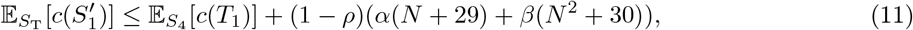

*where the coefficient* (*α*(*N* + 29) + *β*(*N* ^2^ + 30)) *is determined by the maximum values of duplication [7] and deep coalescence [31] scores, respectively*.

We note that the arguments used to derive these inequalities do not explicitly depend on the topology, and hence analogous bounds can be derived for an embedded symmetric structure and an alternative asymmetric structure. Hence, by leveraging this result, we can extend our observation to all trees with 4 or more taxa, proving that no linear combination of the duplication and deep coalescence costs (and by Observation 1 no linear combination of duplication, loss, and deep coalescence costs) is statistically consistent under the MSC model.

#### Theorem 1.

*For any choice of weights α, β* ∈ ℝ *the gene tree parsimony estimator with the cost function c*(*G, S*) = *αc*_*D*_(*G, S*) + *βc*_*X*_ (*G, S*) *is statistically inconsistent for species trees with N* ≥ 4 *taxa under the MSC model*.

*Proof*. Combining the two inequalities from Lemma 4 it follows that the estimator is inconsistent as long as there exist *ρ* satisfying

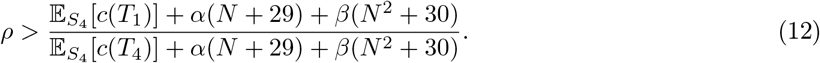

Now, from Lemma 3 we know that the right-hand side of the inequality above is less than 1 for *α/β* < 7 the appropriate choice of parameters, and hence there exists *ρ* satisfying the above condition. Similarly, for *α/β* ≥ 7, we need to find *ρ* that is greater than 1*/*RHS in inequality 12, which is also satisfiable for appropriate parameter choice. Thus, we conclude that for any choice of weights *α, β* the gene tree parsimony estimator with the cost function *c*(*G, S*) = *αc*_*D*_(*G, S*) + *βc*_*X*_ (*G, S*) is statistically inconsistent for species trees with *N* ≥ 4 taxa under the MSC model.

*Remark 1*. The result above shows that given any linear combination of the scores, there exists an anomaly zone in which this combination is inconsistent. However, this does not prove that there exists an anomaly zone that simultaneously misleads both duplication and deep coalescence GTPs evaluated independently. While we do not have a rigorous proof of existence of such an anomaly zone, empirical evidence for it can be provided by a tree that simultaneously exhibits symmetric and asymmetric gadgets (Fig. 2).

**Fig. 2.**
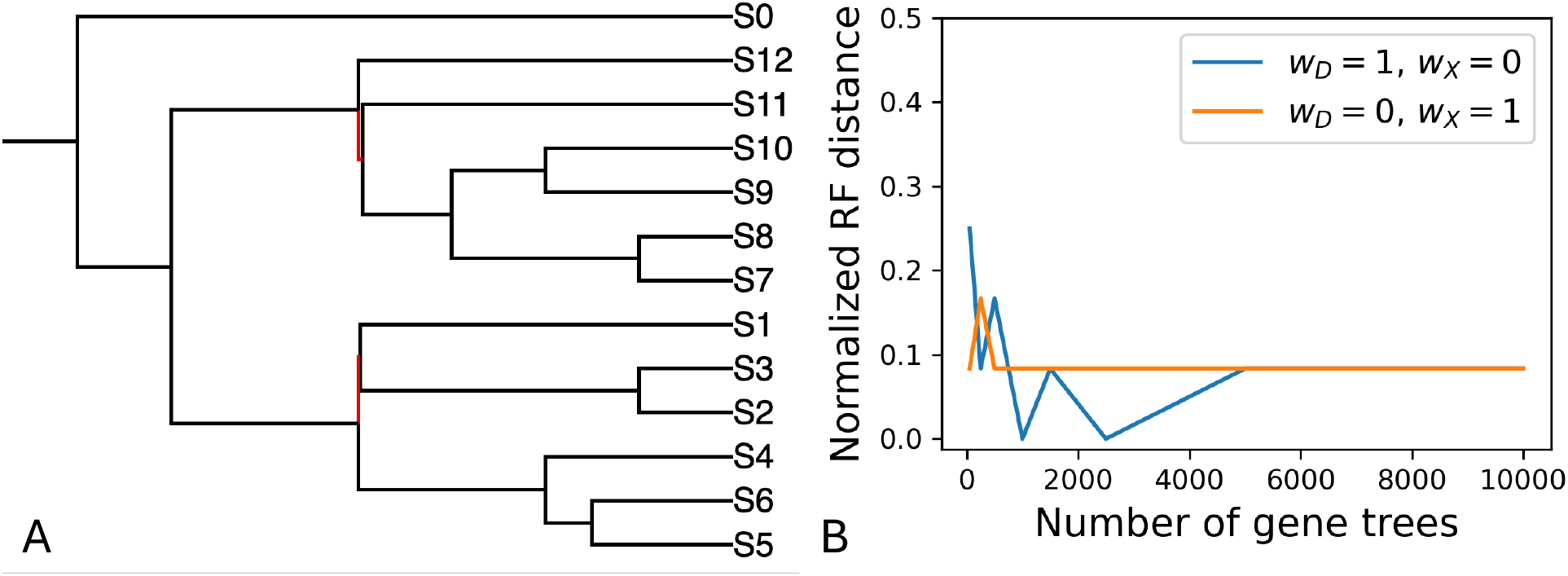
(A) A phylogenetic tree exhibiting both kinds of problematic branches for GTP duplication and deep coalescence costs (highlighted in red). (B) Normalized Robinson-Foulds distance to the ground truth species tree as the function of the number of input gene trees. Neither the duplication-based nor the deep-coalescence-based GTP approach converges to the correct topology. Each approach fails to recover its corresponding anomaly branch.

## 4 Empirical results

### 4.1 Experimental setup

In order to provide an empirical evaluation of GTP methods under MSC with gene duplication and loss, we have designed a set of simulated data experiments, largely following the setup from a prior study by Zhang *et al*. [33] and utilizing SimPhy [15]. We have considered a total of four simulation scenarios: (A) high ILS, duplication and loss rates; (B) low ILS, high duplication and loss rates; (C) low ILS, high duplication, and low loss rates; and (D) high ILS, and high duplication rates, and no loss rate (Table 2).

**Table 2.** Details of the simulation scenarios considered in this study.

| Scenario | Eff. pop. size | Duplication rate | Loss rate |
| --- | --- | --- | --- |
| A | $4.7 \times 10^8$ | $4.9 \times 10^{-10}$ | $4.9 \times 10^{-10}$ |
| B | $4.8 \times 10^7$ | $4.9 \times 10^{-10}$ | $4.9 \times 10^{-10}$ |
| C | $4.8 \times 10^7$ | $4.9 \times 10^{-10}$ | $4.9 \times 10^{-11}$ |
| D | $4.7 \times 10^8$ | $4.9 \times 10^{-11}$ | 0 |

We controlled the level of ILS via the effective population size parameter, and duplication and loss rates via respective duplication/loss rate parameters. For all scenarios, the number of independent replicates was kept at 50. The random seed was kept fixed at 1441 for all simulation scenarios. For scenario A, we simulated species trees with 10, 20, and 50 taxa. For all of the other scenarios, we only considered the 50-taxon case.

#### Species tree simulation

Species trees were simulated with SimPhy [15] under the pure birth model. The birth rate (-sb) was set to 5 × 10^−9^ for all simulations. Tree height (-st) was sampled from the log-normal distribution with the location parameter set to 21.25 and scale parameter set to 0.2. The ratio of ingroup tree height to the length of the branch to the ingroup (-so) was kept at 1 for all simulations. Tree-wide substitution rate (-su) was sampled from the log-normal distribution with the location parameter set to −21.9 and scale parameter set to 0.1. Finally, tree-wide effective population size (-sp) was set to 4.7 × 10^8^ for scenarios A and D, and to 4.8 × 10^7^ for scenarios B and C.

#### Gene tree simulation

For all scenarios 250 and 500 gene trees were simulated with SimPhy [15] with duplication and loss rates specified as indicated in Table 2. Substitution rate heterogeneity was modeled with: species-specific branch rate heterogeneity modifier (-hs) sampled from log-normal distribution (1.5, 1), gene-family-specific rate heterogeneity modifier (-hl) sampled from log-normal distribution (1.551533, 0.6931472), and gene-by-lineage-specific rate heterogeneity modifier (-hg) sampled from log-normal distribution (1.5, 1).

#### Sequence simulation

For each of the four scenarios A, B, C, D we simulated multiple sequence alignments with INDELible [6] for the *N* = 50 case. We simulated three alignments with sequence lengths of 100, 250, and 500 respectively. For all simulated alignments, the same parameters have been used. Substitutions were modeled with the GTR model with rates sampled from a 6-dimensional Dirichlet distribution with parameters (16, 3, 5, 5, 6, 15). Equilibrium probabilities were sampled from a 4-dimensional Dirichlet distribution with parameters (36, 26, 28, 32). Indels were modeled using Zipfian distribution with parameter sampled uniformly from [1.5, 2] and a maximum indel size of 10. The insertion rate was set equal to the deletion rate and sampled uniformly from [0.001, 0.002].

#### Gene tree inference

For the simulated alignments, gene trees were inferred with IQ-TREE [17] under the GTR + *Γ* substitution model with all other parameters set to their default values.

#### Species tree inference

Due to Observation 1 we only focused on weights for duplication and deep coalescence, and furthermore since *c*_*D*_(*G, S*) ≤ *c*_*X*_ (*G, S*) for any *G* and *S* (Proof of Theorem 3.2 [35]), we only evaluated the cases where *w*_*D*_ ≥ *w*_*X*_ . We used DynaDup v2.3.2 [3,28] to infer species tree from the collection of gene trees. We inferred species trees with all possible subsets of cost functions, as well as with duplication cost having weights [2, 4, 8, 16, 32, 64] while the deep coalescence cost had a constant weight 1.

### 4.2 Characteristics of the simulated data

First, we quantified the gene tree estimation error (GTEE) measured as the normalized sum of the false positive and false negative splits in the inferred trees (Fig. 3). As expected, the GTEE decreased for longer alignments for all scenarios. The distribution of GTEE was highly similar among the four scenarios considered, with scenarios A and B showing slightly multimodal behavior of the error, absent from the scenarios C and D (Fig. 3).

**Fig. 3.**
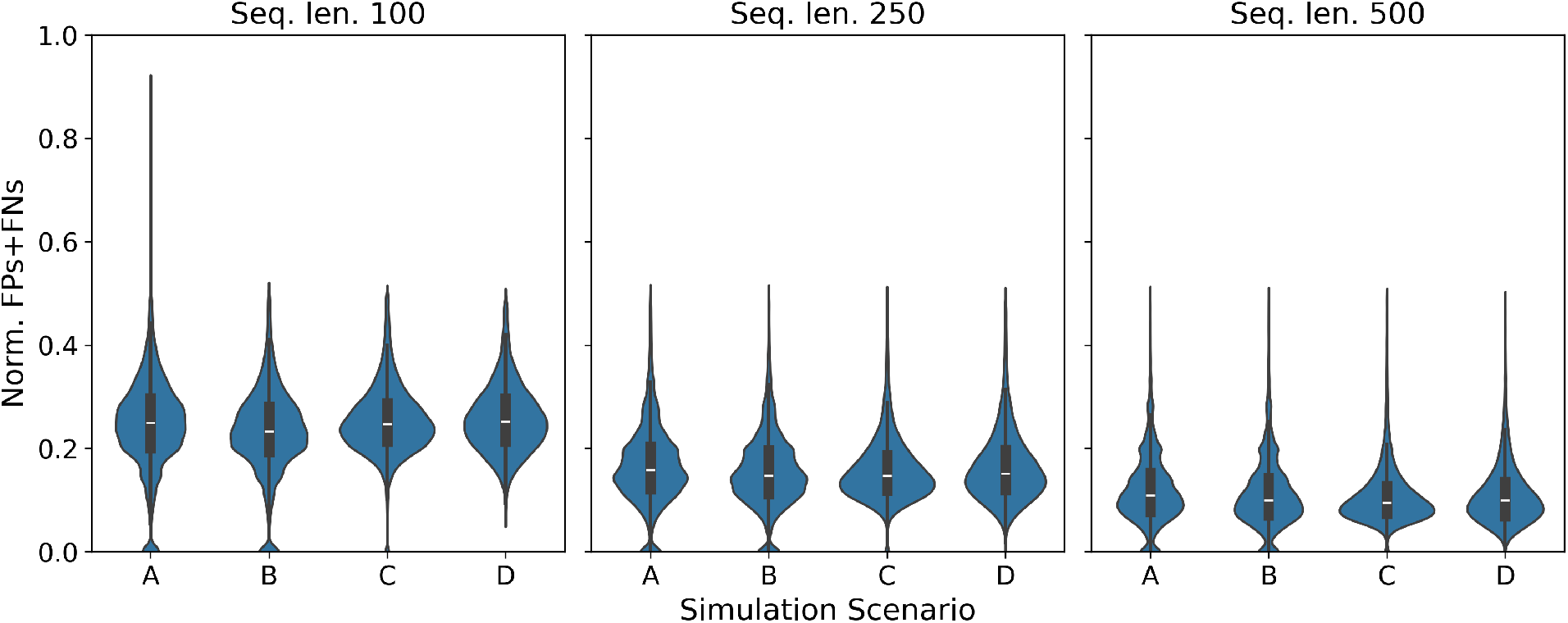
Violin plots showing the distribution of the normalized sum of false positive (FP) and false negative (FN) splits in each of the inferred gene trees. Panels (left to right) show different sequence lengths for the DNA sequence in the simulation. x-axis indicates the simulation scenario. All reported data is for the trees on 50 taxa.

### 4.3 Performance on the simulated gene trees

Next, we have evaluated how GTP methods based on duplication, loss, and deep coalescence costs, and an equal weight linear combination of all three costs perform on the simulated gene tree data in the presence of ILS, gene duplication and loss (Scenario A, Fig. 4). We also included ASTRAL-Pro 3, a method designed for handling paralogs while performing inference under MSC model, as a baseline in the evaluation.

**Fig. 4.**
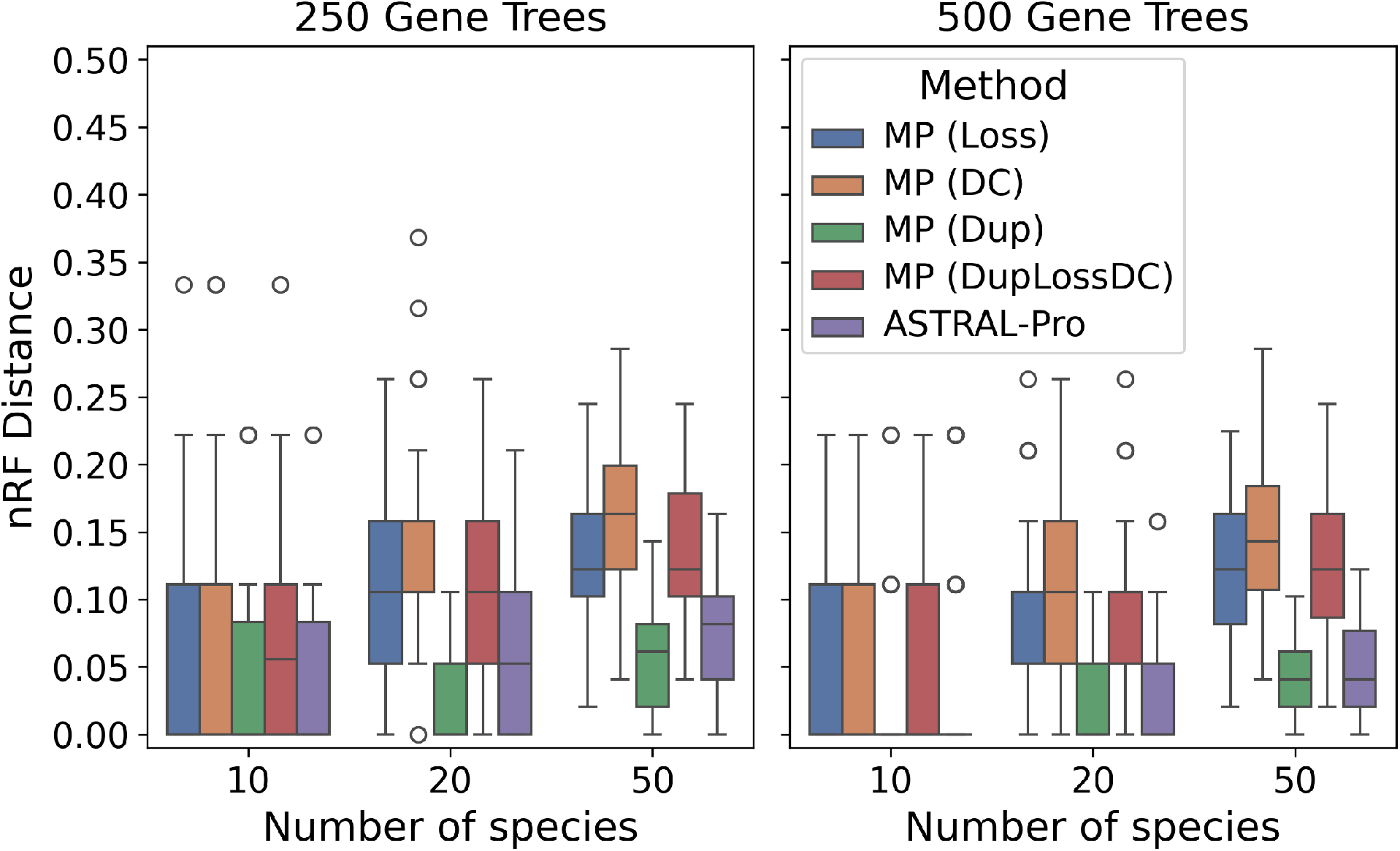
Normalized Robinson-Foulds distance between inferred and true species trees across GTP scoring schemes in scenario. **A**. y-axis shows normalized Robinson-Foulds distance between the true species tree and the one inferred from true gene trees by each of the methods (shown with different colors). x-axis shows the number of species in each instance of scenario A. All boxplots are based on 50 replicate runs.

We observe that as the number of species increases, the topological error in the inferred species trees increases for all methods (Fig. 4). With an increase in the number of gene trees used for inference, the topological error in the inferred species trees goes down for ASTRAL-Pro 3. However, for all GTP methods, the topological error does not consistently decrease as more gene trees are available, with the exception of duplication-based parsimony cost for 10 and 50 species cases (Fig. 4).

Next, we explored how these methods perform under varying duplication and loss rates, as well as at different levels of ILS.

In particular, across the four scenarios considered (Table 2) we observe that all methods have higher error when the ILS levels are higher (Scenarios A, D, Fig. 5). Notably, in all cases GTP method using only the duplication cost performed the best among GTP methods, and achieved topological error values comparable to or better than ASTRAL-Pro 3 (Fig. 5).

**Fig. 5.**
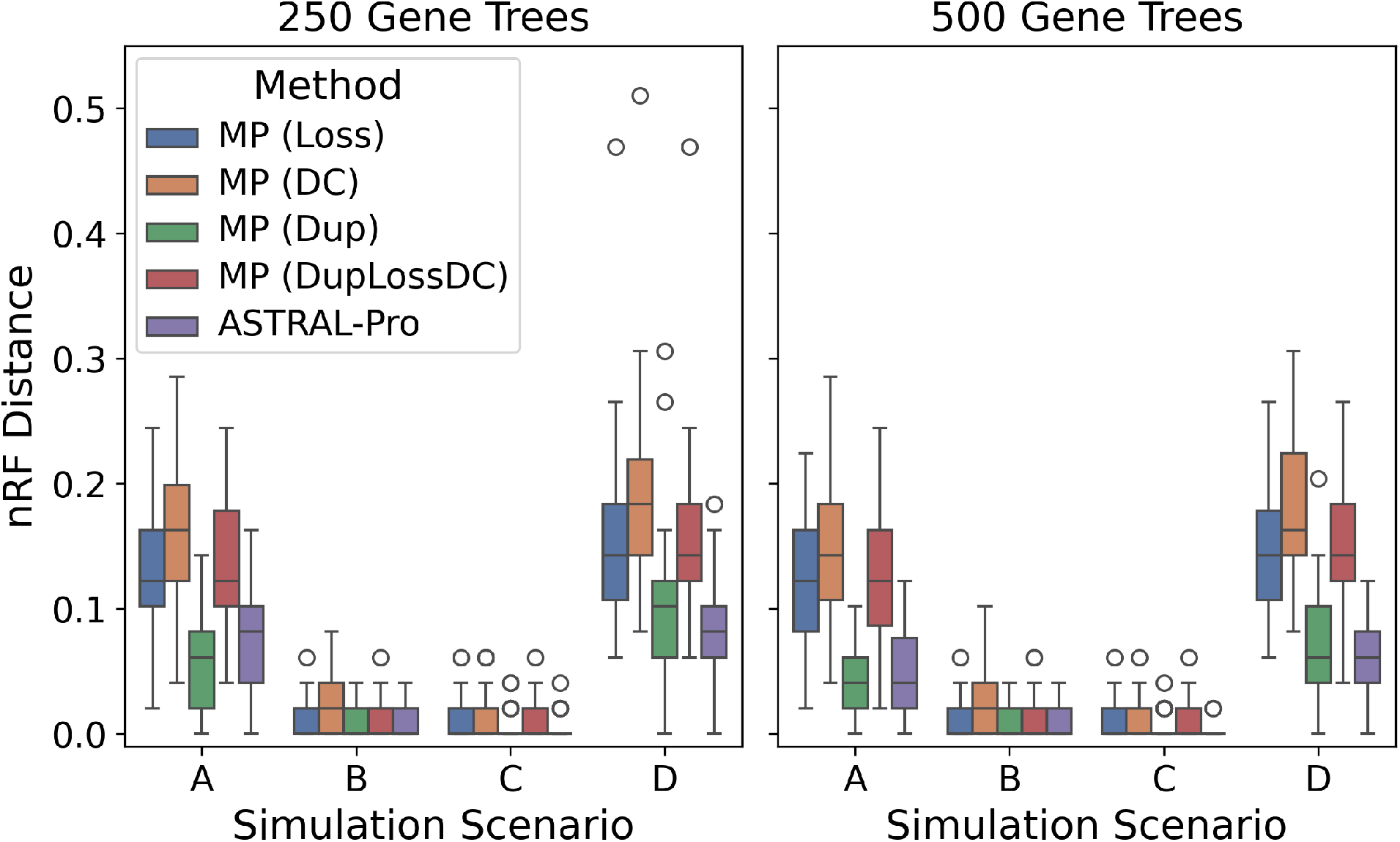
Normalized Robinson-Foulds distance between inferred and true species trees for scenarios. **A-D**. y-axis shows normalized Robinson-Foulds distance between the true species tree and the one inferred from true gene trees by each of the methods (shown with different colors). x-axis shows the simulation scenario defined by the tree-wide effective population size, and duplication and loss rates. All data shown are based on species trees with 50 taxa. All boxplots are based on 50 replicate runs.

To further explore the impact of various weight schemes when combining different GTP scores, we have evaluated how the ratio of the weight associated with the duplication cost (*α*) to the weight associated with the deep coalescence cost (*β*) impacts inference accuracy (Fig. 6).

**Fig. 6.**
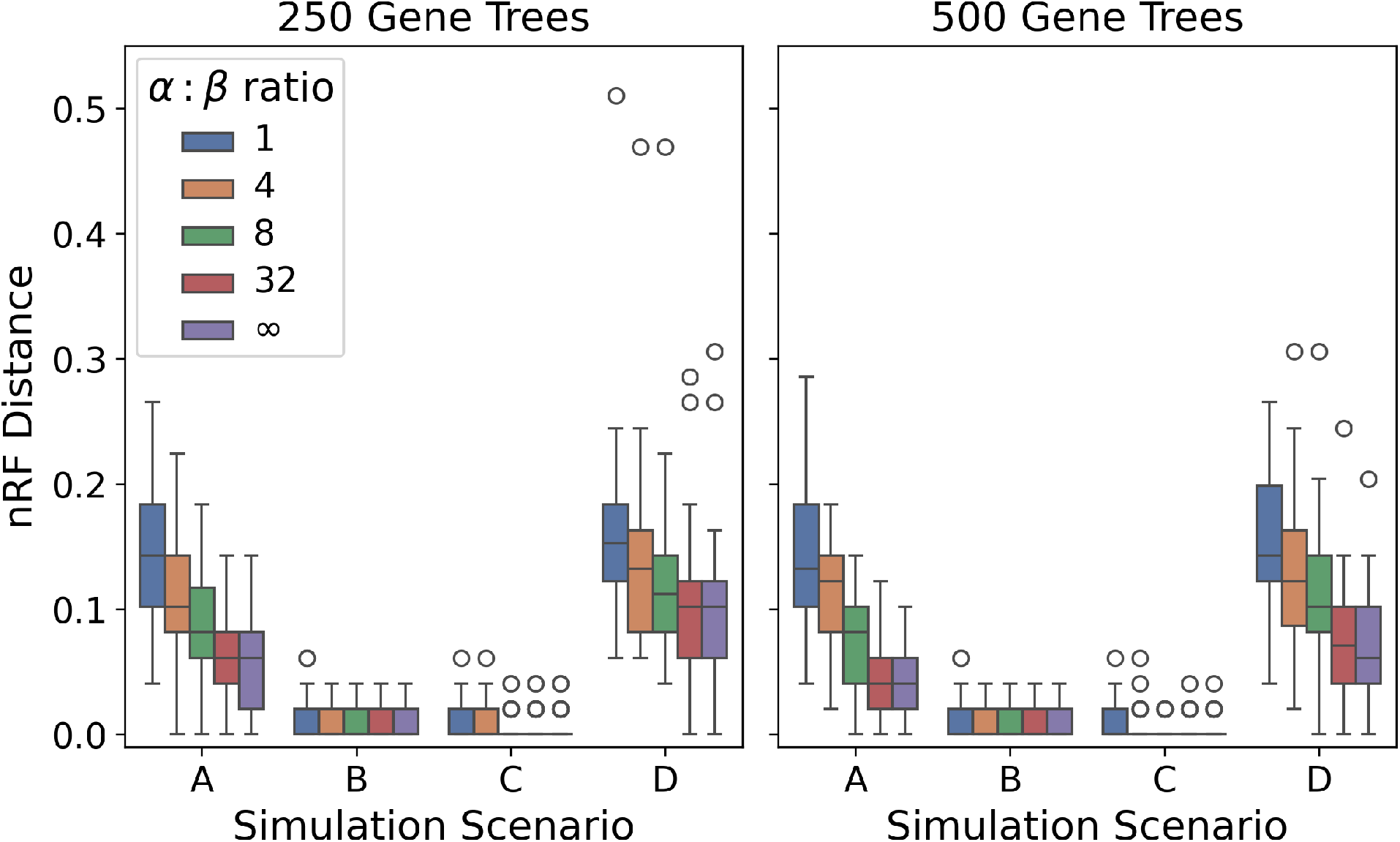
Normalized Robinson-Foulds distance between inferred and true species trees with varying duplication-to-deep-coalescence cost ratios. y-axis shows normalized Robinson-Foulds distance between the true species tree and the one inferred from true gene trees under different ratios of duplication (*α*) to deep coalescence (*β*) costs shown with different colors. x-axis shows the simulation scenario defined by the tree-wide effective population size, and duplication and loss rates. All data shown are based on species trees with 50 taxa. All boxplots are based on 50 replicate runs.

We note that in all cases, as the weight given to duplication cost increases the topological error of the inferred species tree decreases (Fig. 6). This behavior is consistent across all scenarios, with the ratio of 32 achieving performance comparable to the use of duplication cost only (Fig. 6).

### 4.4 Performance on the inferred gene trees

To further evaluate how GTP methods perform in practice, we have evaluated topological accuracy of the inferred species trees in the case when gene trees are inferred from the simulated multiple sequence alignment data (Figs. 7,8). We note that similarly to the experiments on simulated gene trees, ASTRAL-Pro 3 shows an improvement in performance across all scenarios with the increase in the number of gene trees available (Fig. 7, top vs bottom panels). Conversely, no such trend is observed for the GTP methods. Also, similarly to the case of simulated gene trees, in all scenarios we note that minimizing only the duplication cost (Fig. 7, green; Fig. 8) provides the highest accuracy among the GTP scoring schemes. Additionally, the ILS level is the biggest contributor to GTP errors leading to scenarios A and D having the highest topological distances to ground truth species trees.

**Fig. 7.**
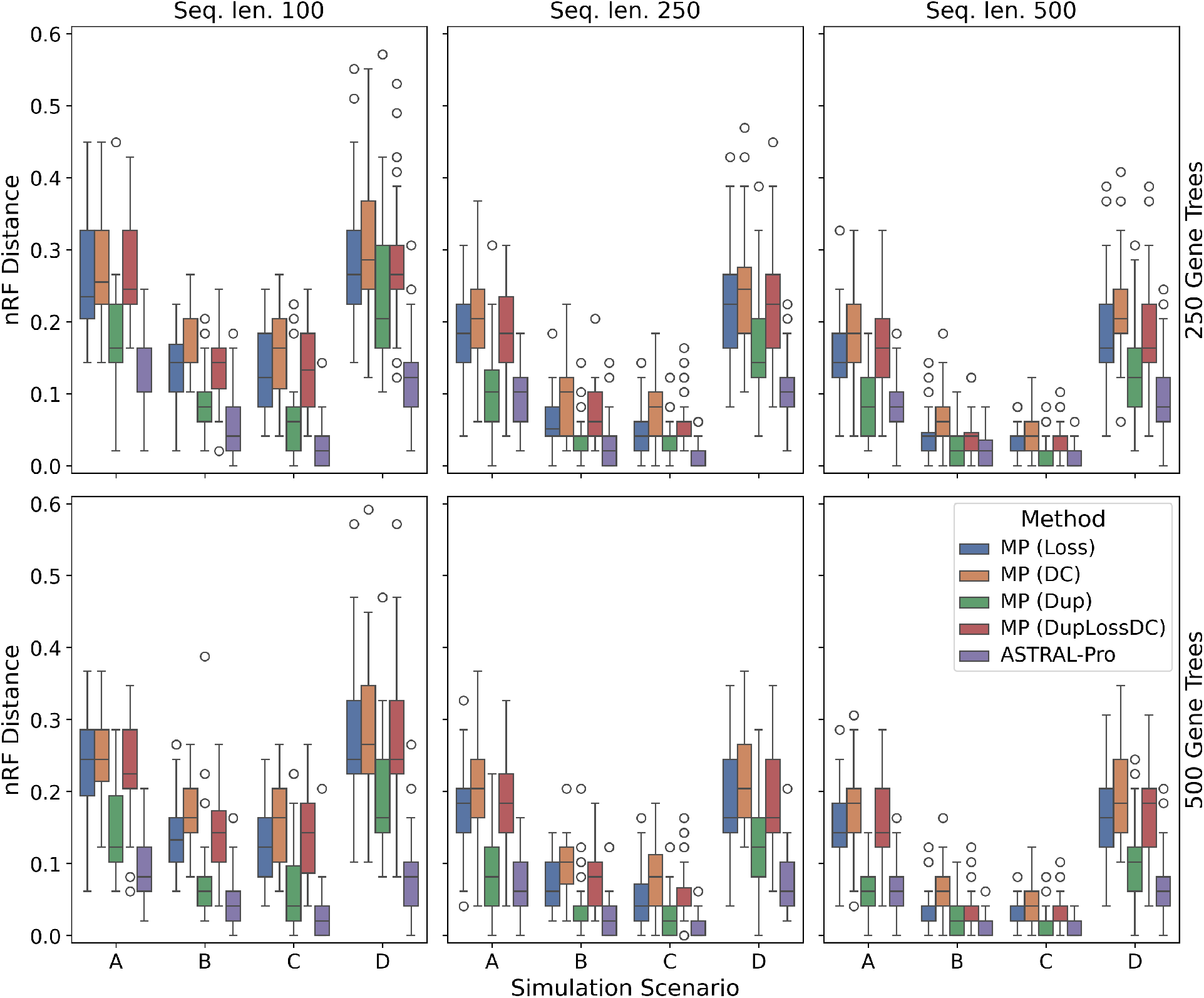
Normalized Robinson-Foulds distance between inferred and true species trees for scenarios. **A-D**. y-axis shows normalized Robinson-Foulds distance between the true species tree and the one inferred by each of the methods (shown with different colors). x-axis shows the simulation scenario defined by the tree-wide effective population size, and duplication and loss rates. All data shown are based on species trees with 50 taxa. Two rows correspond to 250 and 500 inferred gene trees, and each column corresponds to 100, 250, and 500 base pairs in the DNA sequence. All boxplots are based on 50 replicate runs.

**Fig. 8.**
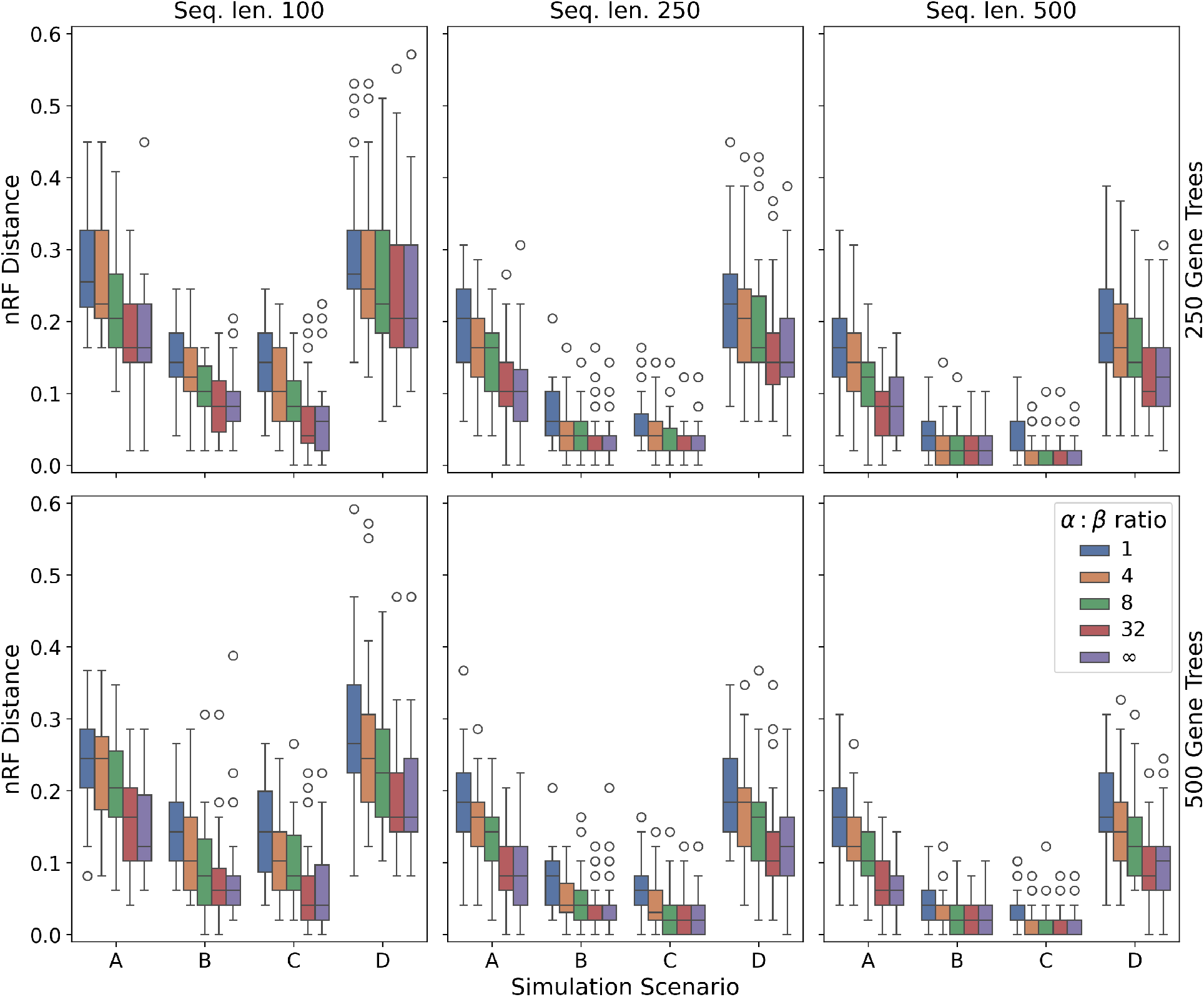
Normalized Robinson-Foulds distance between inferred and true species trees with varying duplication-to-deep-coalescence cost ratios. y-axis shows normalized Robinson-Foulds distance between the true species tree and the one inferred under different ratios of duplication (*α*) to deep coalescence (*β*) costs shown with different colors. x-axis shows the simulation scenario defined by the tree-wide effective population size, and duplication and loss rates. All data shown are based on species trees with 50 taxa. Two rows correspond to 250 and 500 inferred gene trees, and each column corresponds to 100, 250, and 500 base pairs in the DNA sequence. All boxplots are based on 50 replicate runs.

The analysis of the varying ratios of duplication to deep coalescence costs on the inferred gene trees shows the same trends as the one performed on simulated gene trees (Fig. 8). We note that the higher weight given to duplication cost leads to better inference accuracy in all scenarios. Interestingly, unlike in the case of simulated gene trees, in scenario D using the ratio of weights equal to 32 yields better performance than using just the duplication costs alone (Fig. 8, middle and left panels).

### 4.5 Biological fungi data

Finally, we also evaluated performance of the GTP methods on a biological dataset of 16 species of fungi examined in [4,27]. We inferred species trees under various cost schemes for GTP, as well as a single ASTRAL-Pro 3 tree (Fig. 9). All inferred species trees had an identical topology (Fig. 9B) that differed by one split from the topology reported in the prior studies (Fig. 9A). The differing split has been consistently identified by multiple methods on this data in a prior study [32].

**Fig. 9.**
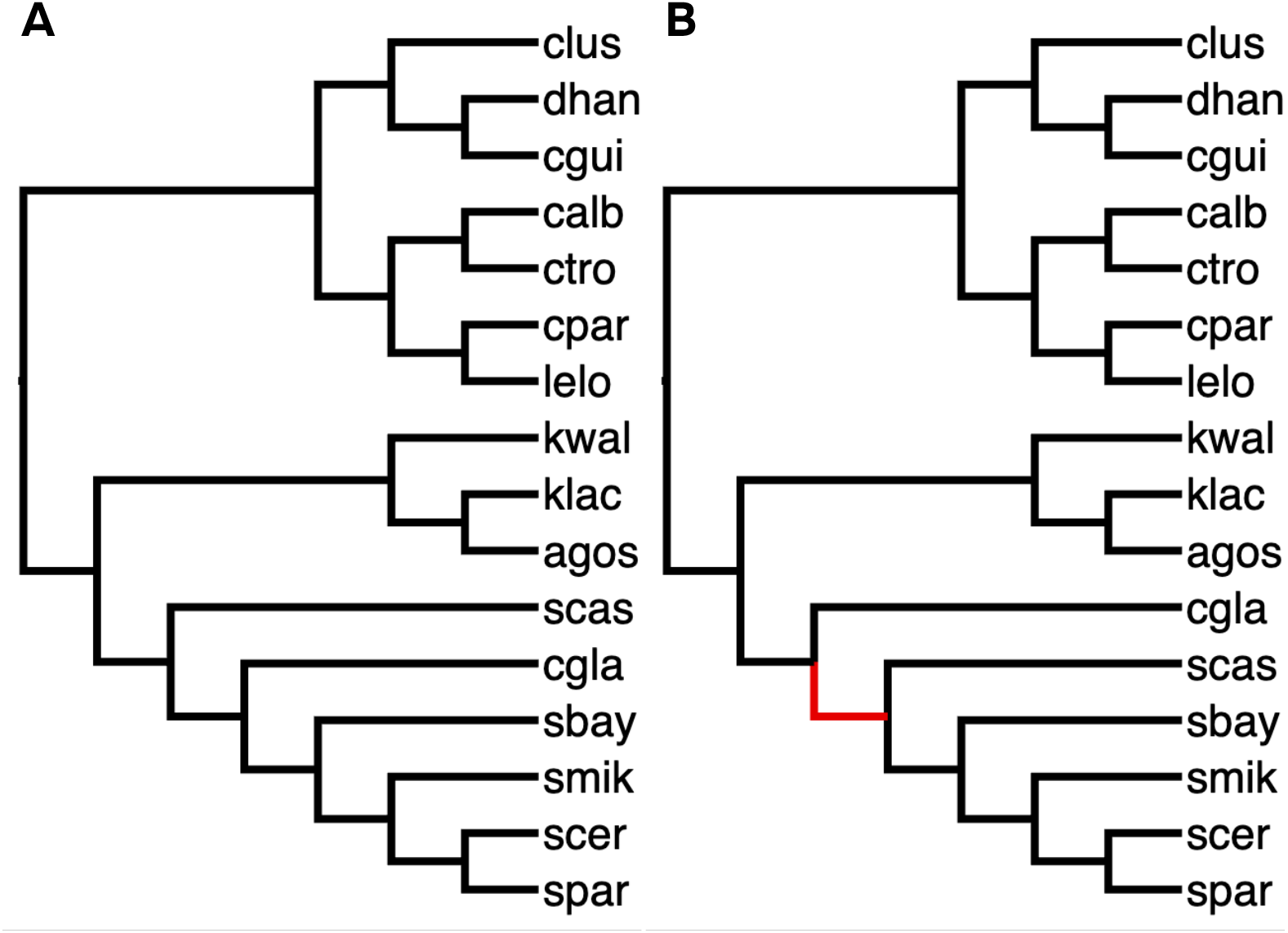
Species tree topologies for 16 species of fungi: (A) the topology reported in the original publications, (B) topology inferred by ASTRAL-Pro 3 and DynaDup. The red edge highlights the single differing split between the topologies (A) and (B).

## 5 Discussion

Theorem 1 proves that no linear combination of gene duplication and deep coalescence costs yields a statistically consistent estimator. However, our empirical evaluation indicates that in the setting when the ILS rate is low, parsimony estimators tend to perform well. This is also supported by work that investigated performance of parsimony methods in the absence of ILS [24]. Furthermore, we note that both theoretical results and empirical analyses suggest that the ratio of deep coalescence cost weight to that of duplication cost should be low in order to minimize the anomaly zone. This is in part due to deep coalescence cost always being greater than or equal to the duplication cost for any given gene tree species tree pair [35].

We note that while our analysis focused on the inconsistency due to ILS under the MSC, additional work is required to properly investigate consistency of different approaches under unified duplication-loss-coalescence (DLCoal) model [27] and multilocus-multispecies coalescent (MLMSC) [10]. Recent results show that quartet-based methods are statistically consistent under some formulations of the DLCoal [16,8,23]. However, theoretical and empirical results on sample complexity, as well as the investigation of certain practical concerns, such as the impact of rooting errors, remain open questions [23].

## 6 Code and data availability

All code used to generate and process the data, as well as to perform all plotting, is available on GitHub: https://github.com/nsapoval/gt-parsimony.

## Acknowledgments

The authors would like to thank Zhi Yan for her feedback on the manuscript. This work was in part supported by the NSF grants DMS/NIGMS-2153704 and DBI-2030604.

